# No evidence for *Wolbachia* as a nutritional co-obligate endosymbiont in the aphid *Pentalonia nigronervosa*

**DOI:** 10.1101/609511

**Authors:** Alejandro Manzano-Marín

**Affiliations:** UMR 1062 Centre de Biologie pour la Gestion des Populations, INRA, CIRAD, IRD, Montpellier SupAgro, Univ. Montpellier, Montpellier, France

**Keywords:** aphid symbiont, co-obligate symbiosis, *Pentalonia nigronervosa*, *Buchnera*, *Wolbachia*

## Abstract

Obligate symbiotic associations are present in a wide variety of animals with a nutrient-restricted diet. Aphids (hemiptera: Aphididae) almost-universally host *Buchnera aphidicola* bacteria in a specialised organs (called bacteriomes). These bacteria supply the aphid with essential nutrients lacking from their diet (i.e. essential amino acids and some B vitamins). Some aphid lineages, such as species from the Lacninae subfamily, have evolved co-obligate associations with secondary endosymbionts, deriving from a loss of biotin-and riboflavin-biosynthetic genes. In this study I re-analyse previously published sequencing data from the banana aphid *Pentalonia nigronervosa*. I show that the metabolic inference results from De Clerck *et al.* (2015) are incorrect and possibly arise from the use of inadequate methods. Additionally, I discuss how the biased interpretation of their antibiotic treatment analyses together with the incorrect metabolic inference resulted in the erroneous suggestion “that a co-obligatory symbiosis between *B. aphidicola* and *Wolbachia* occurs in the banana aphid”.

---

In a previous study, De Clerck *et al.* (2015) claimed to present evidence for a potential co-obligate association of *Buchnera* and *Wolbachia* in the aphid *Pentalonia nigronervosa*. They reach these conclusions mainly based on 4 lines of evidence: **(1)** The apparently fixed nature of *Wolbachia* in *P. nigronervosa*; **(2)** a genome-based metabolic inference coming from a pooled metagenomic assembly of extracted DNA from three *P. nigronervosa* populations sampled in Gabon, Madagascar, and Burundi; **(3)** an antibiotic treatment directed towards the elimination of the endosymbionts; and **(4)** the attempted detection of putatively missing genes through PCR. In this work I have re-analysed the publicly available sequencing data and performed a genome-based metabolic inference of the biosynthetic capabilities of *Buchnera*. I find that this *Buchnera* has equivalent biosynthetic capabilities; concerning essential amino acids, B-vitamins, and co-factors; to all published *Buchnera* strains from “mono-symbiotic” aphids (only harbouring *Buchnera* as the nutritional co-obligate symbiont), and thus should not need an additional partner to fulfil its nutritional role. I also critically discuss the interpretation of the experimental results presented by De Clerck *et al.* (2015) and conclude that their suggestion of the nutritional-based co-obligate nature of *Wolbachia* in *P. nigronervosa* derives from inadequate analyses of their data as well as a biased interpretation of their experimental results.

To produce a genome assembly, I downloaded the three datasets deposited in NCBI with project number PRJNA268300 and accession SRX766492. The pooled genome assembly of these reads resulted in 269,717 contigs that were then binned into 4 groups: *Wolbachia, Buchnera*, mitochondrion, and the aphid host. These resulted in 135 scaffolds for *Buchnera* with an average k-mer (99 bps) coverage of 110 and 1309 for *Wolbachia* with an average k-mer coverage of 396. The search for the genes involved in the biosynthesis of essential amino acids (EAAs), B vitamins and other co-factors (hereafter referred to collectively as “nutritional genes”), revealed that *Buchnera* from *P. nigronervosa* retains all genes common to other *Buchnera* from aphids displaying a mono-symbiotic relationship with *Buchnera* (fig. 1). Additionally, I found that all genes claimed by De Clerck *et al.* (2015) to be missing from *Buchnera* in the biosynthetic pathways shown in figure 4 of the article are actually present, except for that of *pgm* (coding for a phosphoglucomutase which is absent in all currently-sequenced *Buchnera* strains). The *lipA, fabB*, and *bioA* genes show frameshifts in low complexity regions (see GenBank file for details), which is not uncommon for *Buchnera* nor for other small A+T-biased genomes. The expression of these genes is likely to be rescued by ribosomal frameshifting. Additionally, the *trpG* gene also displays a frameshift in a low complexity region and a stop codon in the consensus sequence. Closer inspection revealed that the “TAG” stop codon shows a variant (”CAG”) present at 13.40% in library SRR1662246. This could be explained by the collapsed assembly of the tandem *trpEG* units found in other Aphidinae, which tend to show pseudogenised variants (Baumann *et al.*, 1997; Lai *et al.*, 1996). To test for the presence of the genes claimed as missing by De Clerck *et al.* (2015) and “nutritional genes”, and in a similar fashion to De Clerck *et al.* (2015), I performed read mapping of each library *vs.* the nucleotide sequence of each gene. This confirmed the presence of most of these genes in all three sequencing libraries (supplementary table S1 in supplementary file S1, Supplementary Material online). Many *Buchnera* genes had very low coverages of *≤* 3 in sequencing library SRR1661114 and *≤* 15 in sequencing library SRR1662249. In fact, these two libraries had between 1 - 2% of the reads coming form *Buchnera*, contrasting library SRR1662249, where 25% of the reads mapped to the *Buchnera* scaffold bin (supplementary table S2 in supplementary file S1, Supplementary Material online). The fact that the authors solely searched for intact protein-coding genes, using myRAST (Aziz *et al.*, 2008; Meyer *et al.*, 2008), and their binning method based on a **BLASTX** search *vs.* the nr database of NCBI, rather than a narrower database consisting of expected associates, surely impacted both accurate binning and gene identification. Therefore, the genome-based metabolic inference results do not in fact support the nutritional need of a co-obligate symbiont in *P. nigronervosa*. The lack of amplification of these genes by De Clerck *et al.* (2015) can be explained by the nucleotide sequence divergence between *Buchnera* harboured by not-so-distantly related aphids. In a closer inspection of the primers *vs.* the assembled references, I noted that designed primers did not matched the expected target sequences. Therefore, the lack of amplification is not surprising and should not have been taken as confirmation for the lack of these genes in *Buchnera*.

**Figure 1.**
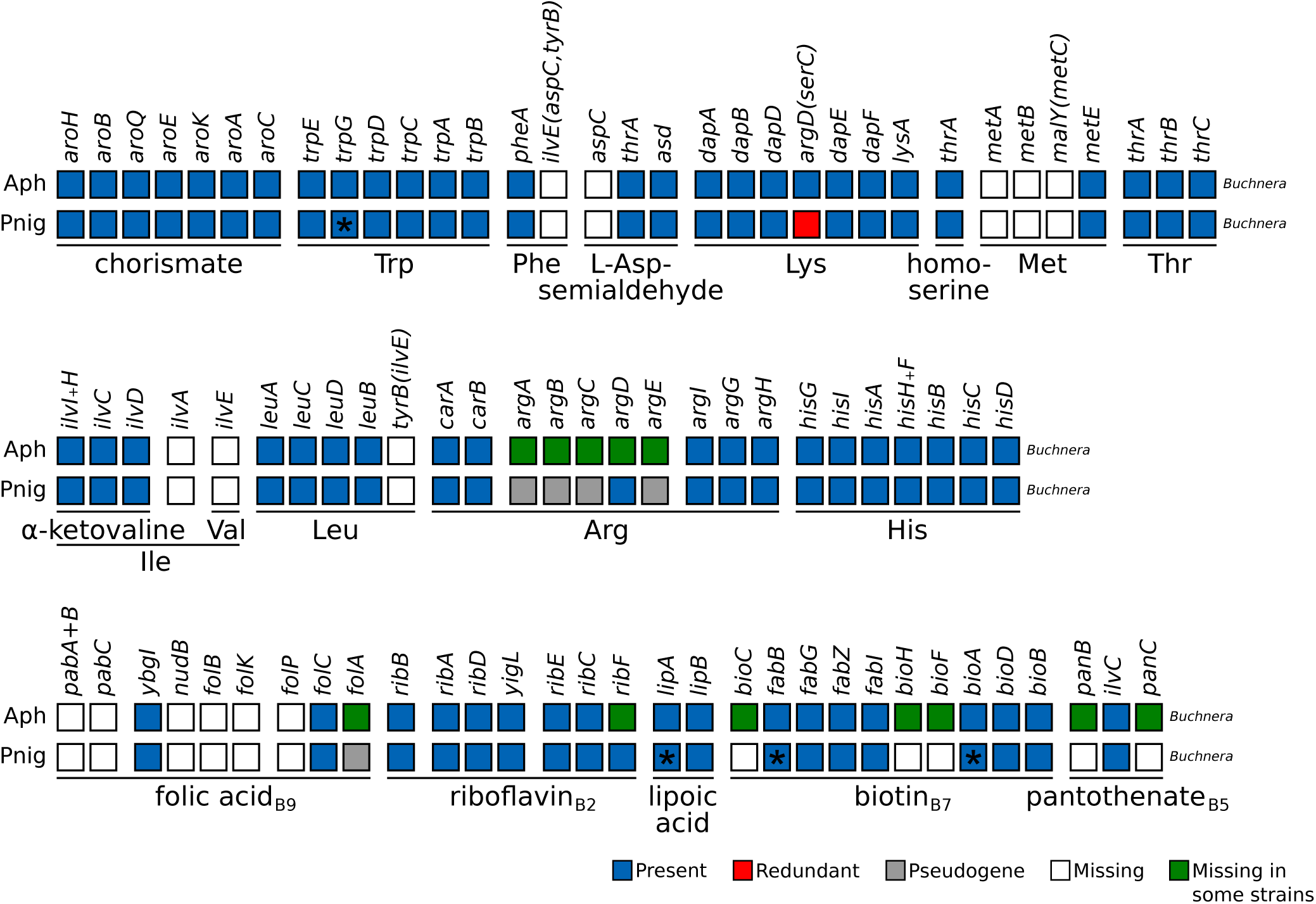
Essential-amino-acid and selected B-vitamin and Cofactor biosynthetic metabolic capabilities of obligate symbiotic consortia of different aphid species. Diagram summarising the metabolic capabilities of *Buchnera* from “mono-symbiotic” Aphidinae aphids (**Aph**) and *P. nigronervosa* (**Pnig**). “Aph” rows are a collapsed representation of several Aphidinae species (see supplementary table S3 in supplementary file S1, Supplementary Material online). The names of genes coding for enzymes involved in the biosynthetic pathway are used as column names. Each row’s boxes represent the genes coded by the *Buchnera* genome. On the bottom, lines underlining the genes involved in the pathway leading to the compound specified by the name underneath the line. For amino acids, its three letter abbreviation is used.

Regarding the experimental analyses, I would argue that in spite of *Wolbachia* being seemingly fixed across different *P. nigronervosa* populations collected in different countries, the fact that this is a facultative endosymbiont cannot be discarded. While it is true that the authors convincingly show that the three populations sampled apparently harbour the same strain of *Wolbachia*, the widespread localisation of this symbiont does not suggest a change of tissue tropism for this strain, but rather what one would expect from a *Wolbachia* bacterium (Ahmed *et al.*, 2015; Gómez-Valero *et al.*, 2004; Schneider *et al.*, 2018). This greatly contrasts the localisation of the nutritional co-obligate *Wolbachia* strain of the bedbug *Cimex lectularius*, where “the infection is substantially restricted to the bacteriome” (Hosokawa *et al.*, 2010). Finally, the reported “selective” elimination of the symbionts through antibiotic treatment does not in fact point towards the obligatory nature of *Wolbachia*. This could only be achieved either through the actual selective elimination of *Wolbachia* and the subsequent measure of biological parameters *vs.* controls (e.g. untreated and/or injected with distilled water), or through proof that the antibiotics solely decrease the quantity of *Wolbachia*, and do not affect *Buchnera*. The authors base their conclusions on the treatments with two antibiotics: tetracycline and ampicillin. Regarding the former, the authors incorrectly conclude that since all the injected nymphs died before adulthood and dead nymphs were only positive for *Buchnera* in PCR assays, then the antibiotic treatment “eliminated *Wolbachia*, but only decreased *Buchnera* quantity, without eliminating it completely”. However, as the authors themselves mention, “tetracycline has been reported to have negative effect on *Buchnera* as well as on *Wolbachia*”. Therefore, it is reasonable to conclude that the impact of the decrease in *Buchnera* could also explain the observed mortality. However, without quantitative PCR and/or microscopy analyses, the impact of the antibiotic treatment on *Buchnera* cannot be correctly assessed. In regards to the ampicillin treatment, similar concerns arise. The fact that the authors could only detect *Buchnera* but failed to measure the impact on the abundance of this symbiont in the dead nymphs, raises concerns regarding the decrease in *Buchnera* being the responsible for the observed mortality.

In brief, De Clerck *et al.* (2015) provide no evidence that the apparent fixation of *Wolbachia* in the analysed populations of *P. nigronervosa* is due to this symbiont being a nutritional co-obligate in this aphid. Its fixation could nonetheless be explained by other processes, such as some sort of reproductive manipulation, protection against viral infections, or the aphid strains sampled actually belonging to one, or a few, “superclone(s)” originally infected with closely-related *Wolbachia* endosymbionts.

All data including the genome assembly, symbiont bins, gene annotation for *Buchnera*, gene sequences, and read mapping results can be found in https://doi.org/10.5281/zenodo.2640354 (last accessed April 15^th^ 2019).

## Materials and Methods

### Sequencing data collection and genome assembly

The tree sequencing libraries (SRR1661114, SRR1662246, SRR1662249) from De Clerck *et al.* (2015) were downloaded from the NCBI SRA database (accession SRX766492) from BioProject (https://www.ncbi.nlm.nih.gov/bioproject/268300, last accessed April 15^th^ 2019). Inspection of the downloaded fastq files with **fastQC** v0.11.7 (https://www.bioinformatics.babraham.ac.uk/projects/fastqc/, last accessed April 15^th^ 2019) revealed contamination of the reads with TruSeq adapter indexes 4,5, and 6, as well as an unknown T+G-rich adapter (”TGTGTTGGGTGTGTTGGGTGTGTTGGGTGTGTTGGGTGTGTTGGGT-GTGT”). To trim these adapters from the read datasets and perform quality trimming, I used **fastx clipper** followed by **fastx quality trimmer** which performed right-tail quality trimming (using a minimum quality threshold of 20) and reads shorter than 50 bps were dropped (options -t 20 -l 50). Additionally, **PRINSEQ** v0.20.4 (Schmieder and Edwards, 2011) was used to remove reads containing undefined nucleotides as well as separate those left without a pair after the filtering and trimming process. Assembly of the three libraries was done using **SPAdes** v3.10.1 (Bankevich *et al.*, 2012) with k-mers 33, 55, and 77.

### Binning of assembled sequences to *Buchnera* and *Wolbachia*

First, all scaffolds shorter then 200 bps and with an average k-mer coverage lower than 3 were dropped. This resulted in 10,097 (3.74% of the total number of scaffolds) which were then used for a BLASTX-based binning. This was done by building a database of the proteomes of the Pea aphid and a set of *Wolbachia* and *Buchnera* bacteria. A separate database of the mitochondrion proteins encoded by the aphid *Schizaphis graminun* was built to detect mitochondria sequences. All these protein datasets were available before the time of publication of De Clerck *et al.* (2015). Accession numbers and information on strains can be found in supplementary table S4 (supplementary file S1, Supplementary Material online). A BLASTX search against the aforementioned databases was done using the filtered scaffolds as query (-soft masking true -seg yes -max target seqs 10000 -evalue 1e-3) followed by assigning the scaffold to the organism with the best BLASTX hit to each scaffold. These bins were then filtered by setting a minimum k-mer coverage threshold of 10 for both *Buchnera* and *Wolbachia*. This resulted in 135 and 1,309 scaffolds being assigned to each bacterium, respectively.

### Identification of “missing” and “nutritional genes” in *Buchnera*

For verifying the absence of the genes claimed as missing by De Clerck *et al.* (2015) and the “nutritional genes”, I collected the amino acid sequences of these genes from *Buchnera* strain 5A, APS, and Bp. Then, a TBLASTN search of the aforementioned sequences *vs.* both the total filtered scaffolds and the *Buchnera* scaffold bin (-evalue 1e-03 -db gencode 11) was performed. Afterwards, manual verification of each hit was performed using **UGENE** v1.29.0 and then BLASTX on the on-line BLAST server from NCBI *vs.* the nr database. If the gene was not found in the *Buchnera* scaffold bin, then the gene was verified from the TBLASTN *vs.* the filtered scaffolds. This revealed the misassignment of ‘NODE 9’ to the *Wolbachia* bin. This manual curation also revealed remnants of the ’TGTGTTGGGTGTGTTGGGTGTGTTGGGTGTGTTGGGTGTGTTGGGTGTGT’ sequence inside the *lipA* and *pgi* genes. Further manual inspection of the reads revealed a consistent insertion of this sequence in a specific place of each gene, revealing the biased insertion of the contaminant. Additionally, an alignment of each read library *vs.* the above-mentioned *Buchnera* genes from *P. nigronervosa* was done with **Bowtie2** v2.3.4.1 (Langmead and Salzberg, 2012) for visualisation purposes. To avoid reads from *Wolbachia* mapping to sequences of orthologous genes from *Buchnera*, the *Wolbachia* scaffolds were also included in the Bowtie2 mapping.

## Acknowledgements

This work was supported by the *Marie-Curie AgreenSkills+* fellowship programme co-funded by the *EU’s Seventh Framework Programme* (FP7-609398). This publication has been written with the support of the AgreenSkills+ fellowship programme which has received funding from the EU’s Seventh Framework Programme under grant agreement No. FP7-609398 (AgreenSkills+ contract). I am grateful to the genotoul bioinformatics platform Toulouse Midi-Pyrenees (Bioinfo Genotoul) for providing help and/or computing and/or storage resources. The funders had no role in study design, data collection and analysis, decision to publish, or preparation of the manuscript.

## References

Ahmed M. Z, Li S.-J, Xue X, Yin X.-J, Ren S.-X, Jiggins F. M, Greeff J. M, and Qiu B.-L. 2015. The intracellular bacterium *Wolbachia* uses parasitoid wasps as phoretic vectors for efficient horizontal transmission. PLoS pathogens, 10(2): e1004672. URL http://journals.plos.org/plospathogens/article?id=10.1371/journal.ppat.1004672.

Aziz R. K, Bartels D, Best A. A, DeJongh M, Disz T, Edwards R. A, Formsma K, Gerdes S, Glass E. M, Kubal M, Meyer F, Olsen G. J, Olson R, Osterman A. L, Overbeek R. A, McNeil L. K, Paarmann D, Paczian T, Parrello B, Pusch G. D, Reich C, Stevens R, Vassieva O, Vonstein V, Wilke A, and Zagnitko O. 2008. The RAST server: Rapid annotations using subsystems technology. BMC Genomics, 9(1): 75. URL http://bmcgenomics.biomedcentral.com/articles/10.1186/1471-2164-9-75.

Bankevich A, Nurk S, Antipov D, Gurevich A. A, Dvorkin M, Kulikov A. S, Lesin V. M, Nikolenko S. I, Pham S, Prjibelski A. D, Pyshkin A. V, Sirotkin A. V, Vyahhi N, Tesler G, Alekseyev M. A, and Pevzner P. A. 2012. SPAdes: a new genome assembly algorithm and its applications to single-cell sequencing. Journal of computational biology: a journal of computational molecular cell biology, 19(5): 455–477. URL http://online.liebertpub.com/doi/abs/10.1089/cmb.2012.0021.

Baumann L, Clark M. A, Rouhbakhsh D, Baumann P, Moran N. A, and Voegtlin D. J. 1997. Endosymbionts (*Buchnera*) of the aphid *Uroleucon sonchi* contain plasmids with *trpEG* and remnants of *trpE* pseudogenes. Current Microbiology, 35(1): 18–21. URL http://link.springer.com/10.1007/s002849900204.

De Clerck C, Fujiwara A, Joncour P, Léonard S, Félix M.-L, Francis F, Jijakli M. H, Tsuchida T, and Massart S. 2015. A metagenomic approach from aphid’s hemolymph sheds light on the potential roles of co-existing endosymbionts. Microbiome, 3(1): 63. URL http://microbiomejournal.biomedcentral.com/articles/10.1186/s40168-015-0130-5.

Gómez-Valero L, Soriano-Navarro M, Perez-Brocal V, Heddi A, Moya A, Garcia-Verdugo J. M, and Latorre A. 2004. Coexistence of *Wolbachia* with *Buchnera aphidicola* and a secondary symbiont in the aphid *Cinara cedri*. Journal of Bacteriology, 186(19): 6626–6633. URL http://jb.asm.org/cgi/doi/10.1128/JB.186.19.6626-6633.2004.

Hosokawa T, Koga R, Kikuchi Y, Meng X.-Y, and Fukatsu T. 2010. *Wolbachia* as a bacteriocyte-associated nutritional mutualist. Proceedings of the National Academy of Sciences of the United States of America, 107(2): 769–774. URL http://www.pnas.org/content/107/2/769.long.

Lai C. Y, Baumann P, and Moran N. A. 1996. The endosymbiont (*Buchnera*sp.) of the aphid*Diuraphis noxia*contains plasmids consisting of*trpEG*and tandem repeats of*trpEG*pseudogenes. Applied and environmental microbiology, 62(2): 332–339. URL http://www.ncbi.nlm.nih.gov/pubmed/8593038.

Langmead B and Salzberg S. L. 2012. Fast gapped-read alignment with Bowtie 2. Nature Methods, 9(4): 357–359. URL http://dx.doi.org/10.1038/nmeth.1923

Meyer F, Paarmann D, D’Souza M, Olson R, Glass E, Kubal M, Paczian T, Rodriguez A, Stevens R, Wilke A, Wilkening J, and Edwards R. 2008. The metagenomics RAST server – a public resource for the automatic phylogenetic and functional analysis of metagenomes. BMC Bioinformatics, 9(1): 386. URL https://bmcbioinformatics.biomedcentral.com/articles/10.1186/1471-2105-9-386.

Schmieder R and Edwards R. 2011. Quality control and preprocessing of metagenomic datasets. Bioinformatics, 27(6): 863–864. URL http://bioinformatics.oxfordjournals.org/cgi/doi/10.1093/bioinformatics/btr026.

Schneider D. I, Parker A. G, Abd-alla A. M, and Miller W. J. 2018. High-sensitivity detection of cryptic *Wolbachia* in the African tsetse fly (*Glossina* spp.). BMC Microbiology, 18(S1): 140. URL https://bmcmicrobiol.biomedcentral.com/articles/10.1186/s12866-018-1291-8.

